# AutoScreen: AI Co-Scientist System for Target Discovery in Functional Genomics

**DOI:** 10.64898/2026.09.06.749678

**Authors:** Yuanhao Qu, Xufeng Liu, Xiaotong Wang, Meitong Chen, Xiao Luo, Luna Lyu, Ming Yin, Jingwen Hui, Di Yin, Ravi Dinesh, Lin Qiu, Kexin Huang, Hanchen Wang, Shangyuan Tong, Henry Cousins, Ricky Runchuan Feng, Oswaldo Martinez, Junze Zhang, Tanming Chen, Russ Altman, Jure Leskovec, Aviv Regev, Mengdi Wang, Le Cong

## Abstract

Target discovery in functional genomics remains largely manual and time-consuming, lacking systematic tools for efficient and reproducible gene-level hypothesis generation. We introduce *AutoScreen*, an AI co-scientist system supporting target discovery through both *Pre-screen Design*, which constructs perturbation libraries *de novo* from free-text research descriptions, and *Post-screen Analysis*, which re-ranks experimental screen hits by integrating statistical scores with biological context. AutoScreen leverages the complementary strengths of multiple specialist agents that perform deep research into multi-modal evidence, information restructuring, parallel searches across >26 biomedical databases, evidence synthesis, and target review to provide transparent, explainable gene prioritization with pipeline provenance. Across 320 genome-scale CRISPR screens as expert-curated benchmarks, AutoScreen achieved a ∼19% increase in validated-hit recovery among its top 100 predictions, relative to the strongest agent baseline, and required a 1.2-fold smaller library to recover the same number of hits at the top-500 reference point. AutoScreen reached mean average precision more than two orders-of-magnitude above random baseline. Further, we validated AutoScreen in cancer immune-evasion case studies focusing on natural killer (NK) and T-cell therapeutics. AutoScreen recovered NK-resistance genes that were initially lower-ranked in a leukemia screen, moving *MUC1*, *PDPN*, and *LRRC15* from raw ranks of 118, 81, 1384 to 5, 44, 659, respectively. In follow-up tumor killing assay with primary human NK cells, individual perturbation validated all three hits successfully. Next, in prospective benchmarking across two cytotoxic T-cell-killing screens, AutoScreen recovered 77.1% of ground-truth hits called by the consensus of two gold-standard analysis pipelines (FDR<0.1), achieving 8% improvement over the strongest general-purpose LLM agent baseline. Finally, we re-analyzed >500 public datasets to construct the *AutoScreen Resource Hub*, a growing knowledge base of pre-computed, annotated reports for CRISPR screens, RNA-seq differential expression, and gene-level UK Biobank genome-wide association studies. This agentic AI approach enables real-time genomics benchmark construction that are continuously updated, expanded, for evaluating frontier AI co-scientists. Overall, AutoScreen enables AI-powered target discovery to be more auditable, scalable, extensible, and reproducible.

## Introduction

High-throughput functional genomics has dramatically expanded our ability to interrogate gene function at scale. CRISPR-based screens, expression profiling and genome-wide association studies (GWAS) now routinely nominate hundreds of candidate genes across diverse biological contexts.^1–3^ This abundance has shifted the bottleneck from data generation to prioritization: deciding which candidates merit follow-up. The problem arises in two recurring scenarios—selecting a subset of hits for deeper study after an initial experiment and designing focused libraries when the full perturbation space cannot be interrogated, whether because of practical limits in single-gene studies (*in vivo* experiments, scarce primary cells) or because combinatorial designs make the pairwise or higher-order space prohibitively large. In both cases, target selection still depends on time-consuming literature review, specialized domain knowledge and subjective judgement, limiting throughput, novelty, reproducibility and the cumulative comparability of functional genomics studies.

Existing approaches each address part of the problem. Statistical methods such as differential expression or enrichment analysis yield quantitative rankings^4,5^ but lack the biological context needed to interpret them mechanistically. Knowledge-based approaches that draw on pathway databases or literature mining supply context but are typically siloed within a single domain and integrate heterogeneous evidence poorly, forcing biologists to manually reconcile fragmented signals against an ever-expanding literature. Most recently, LLM-driven agents for experimental design—exemplified by systems that iteratively propose small perturbation panels^6^—have shown that language models can encode useful biological priors. In practice, however, these agents tend to primarily recover well-studied genes, scale poorly to genome-wide library sizes, and offer little provenance for why a target was chosen, leaving their recommendations difficult to audit or trust. An effective solution would need to couple the quantitative rigor of statistical scoring with the contextual breadth of knowledge integration, while exposing the evidence behind every recommendation. It would support both *de novo* library design for prospective investigations and (re)analysis of existing datasets, adapt its reasoning to the specific experimental context, and operate at genomic scale.

Here, we introduce AutoScreen, a modular multi-agent system built to meet these requirements. AutoScreen orchestrates five specialist agents—spanning deep literature research, information restructuring, parallel search across 26 specialized databases, context-aware evidence synthesis and target review—to transform free-text research descriptions into transparent, evidence-backed gene rankings with pipeline provenance. Its two operational modes, Pre-screen Design and Post-screen Analysis, support library construction from a study description and context-aware re-ranking of experimental hit lists, respectively. Because the synthesis step accepts interchangeable model backends, the reasoning component can be substituted as language models advance, with backend selection performed offline against experimentally validated targets. We benchmark AutoScreen across 320 genome-scale CRISPR screens, where it recovers substantially more validated hits at reduced library sizes than general-purpose LLM, agent and random baselines. We then apply AutoScreen to two immune-evasion case studies. Post-screen re-ranking recovered genuine NK-resistance hits (MUC1, PDPN, LRRC15) and generalized to a melanoma CRISPRa screen, where AutoScreen prioritizes the CEACAM family (CEACAM1, CEACAM5, CEACAM6). Pre-screen library design, benchmarked prospectively against two independent cytotoxic-T-cell-killing screens that were not available to the agent, recovered 77.1% of ground-truth hits on average, an 8% relative improvement over the general-purpose LLM baseline design (71.4%), while generating each designed library in 35.54 minutes on average versus approximately 3 days for manual expert curation (Fig. 5h). Finally, we release the AutoScreen Resource Hub with pre-computed analyses of over 500 public CRISPR, RNA-seq and GWAS datasets, lowering the barrier to evidence-based target discovery across the community. By turning unstructured biological knowledge into auditable target prioritization, AutoScreen aims to make functional genomics more efficient, reproducible and generative of new hypotheses.

## Results

### A modular multi-agent system for evidence-based target discovery

AutoScreen is a modular, auditable multi-agent system that automates target prioritization for functional genomics (Fig. 1). It combines deep research over text-based evidence, parallel knowledgebase retrieval across multiple domains, context-aware anchored gene emission and agentic reasoning to return transparent, evidence-backed recommendations.

**Figure 1.**
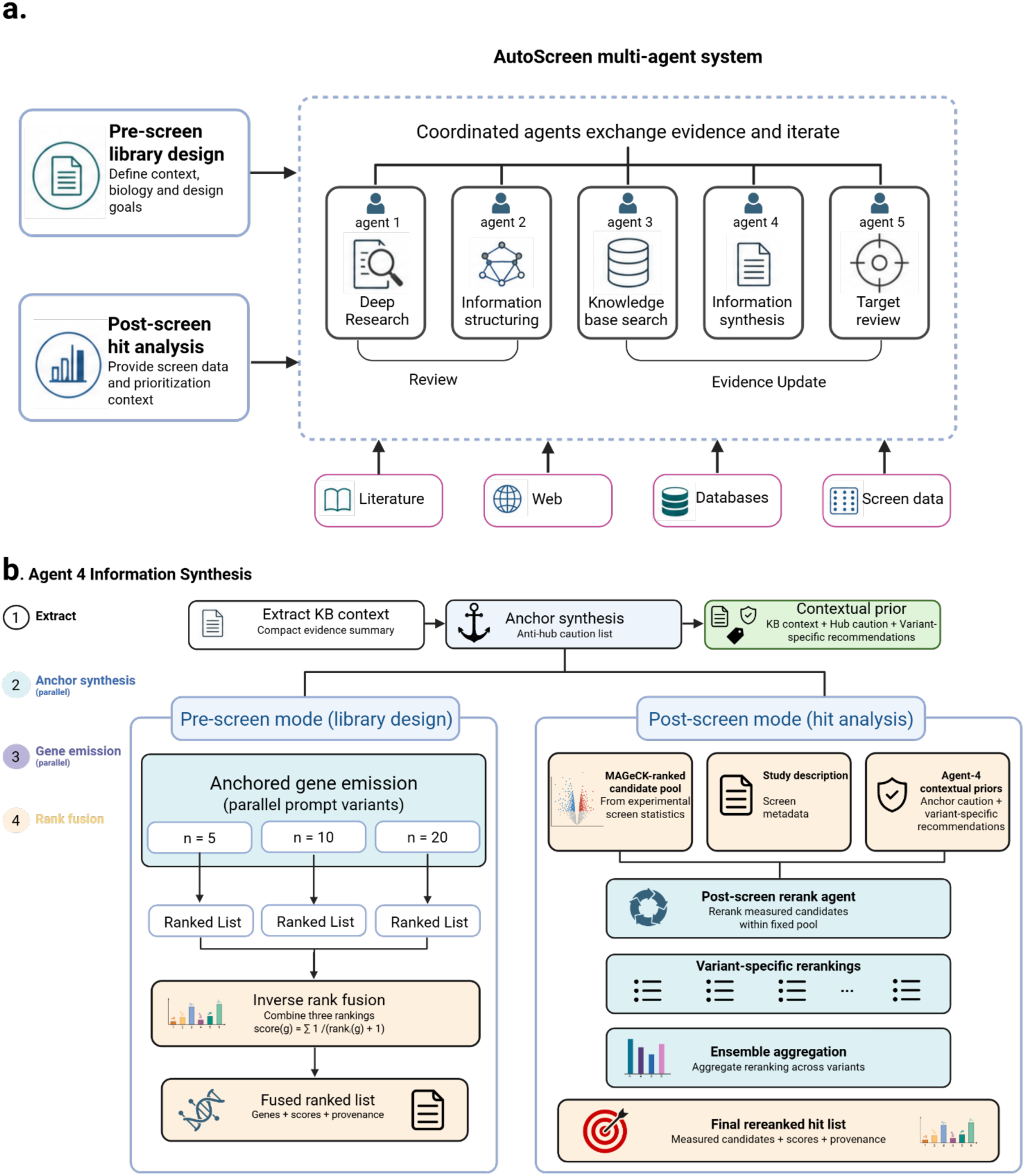
AutoScreen facilitates target discovery in functional genomics through a coordinated multi-agent workflow. **a, System overview.** AutoScreen supports two modes: pre-screen library design from biological context and design goals, and post-screen hit analysis from experimental screen data. Five coordinated agents perform deep research, information structuring, knowledge-based search, information synthesis and target review to generate evidence-backed target reports. **b, Information Synthesis Agent workflow details.** Agent 4 distills knowledge-based outputs into compact biological context and screen-specific anti-hub anchors. In pre-screen mode, it generates ranked candidate-gene lists across multiple prompt settings and fuses them by inverse-rank fusion. In post-screen mode, it passes contextual priors to a re-rank/re-prioritize agent, which re-ranks a MAGeCK-defined candidate pool and aggregates variant-specific rankings into a final hit list with scores and provenance.

The system runs in two modes that address prioritization at different experimental stages (Fig. 1a, left). In Pre-screen Design mode, AutoScreen accepts a free-text study description—for example, “*study the genes that maintain cellular cholesterol levels under protein accumulation in HeLa cells over 18 days*”—and constructs a *de novo* gene perturbation library with full evidence documentation. In Post-screen Analysis mode, it ingests an experimental hit list together with the study description and statistical metrics (*P* values, log fold changes) from tools such as MAGeCK or DESeq2^4,7^, then re-ranks candidates by combining source-level evidence with the original screening signal.

Five specialist agents convert a query into a ranked report (Fig. 1a, right). The Deep Research Agent mines literature, LLM-derived insights, and web knowledge into coherent narrative summaries; the Information Restructure Agent distills these into ontology-aligned search terms; the Knowledge Base Search Agent deploys parallel subagents across over 26 specialized databases spanning pathways, clinical data, interactions, chemicals and drugs; the Information Synthesis Agent converts knowledgebase evidence into screen-specific biological context, synthesizes anti-hub anchors, and fuses anchor-conditioned ranked gene emissions; and the Target Review Agent annotates candidates with expression, dependency and druggability flags before producing the final report. Rather than training a bespoke model, the Information Synthesis Agent accepts interchangeable reasoning backends, which can be selected offline against experimentally validated targets.

### AutoScreen’s Pre-screen Design outperforms general-purpose LLM, agent, and random baselines across 320 expert-curated CRISPR screen benchmarks

To benchmark target discovery at scale, we assembled 320 genome-scale CRISPR screens from BioGRID ORCS^8^ that met stringent inclusion criteria (complete metadata and a condition- or phenotype-specific contrast; Fig. 2a), defining the top 100 genes per screen as ground-truth hits. From each screen’s metadata we curated a research goal—for example, “*study how 2-day interleukin-1β exposure affects A375 tumor resistance to NK killing*.” The benchmark spanned loss-of-function screens using CRISPR knockout and CRISPR interference, addressing drug and chemical responses, viral and bacterial infections, protein accumulation and other cell biological and functional phenotypes. We compared AutoScreen against BioDiscovery Agent (an LLM agent that iteratively designs small perturbation panels), GPT-5.4 (a general-purpose LLM)^9^ and a random baseline. All methods received only the curated research goal and returned a ranked gene list in Pre-screen Design mode.

**Figure 2.**
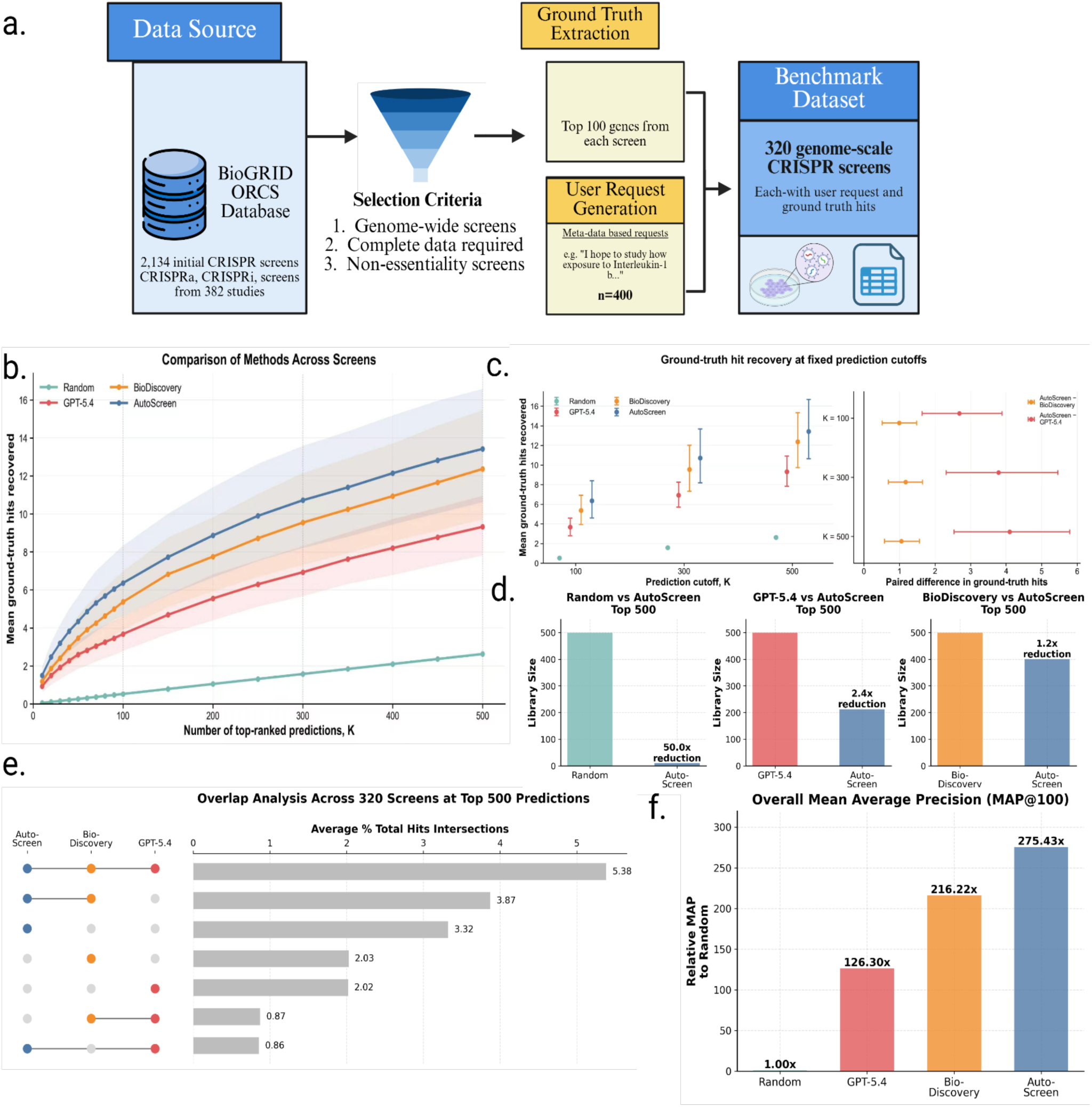
AutoScreen improves target recovery and ranking across genome-scale CRISPR-screen benchmarks. **a,** Benchmark construction from public genome-scale CRISPR screens. Screens meeting inclusion criteria were retained, ground-truth target sets were defined from top-ranked genes in the original screen analyses, and screen metadata were converted into user-style target-discovery queries. **b,** Average recovery of ground-truth hits as a function of prediction-list size for AutoScreen, BioDiscovery, GPT-5.4 and random selection. Lines and points indicate mean recovery across screens; shaded bands indicate 95% confidence intervals. Curves are evaluated at prediction cutoffs of 10–100 in increments of 10 and 150–500 in increments of 50. **c,** Ground-truth hit recovery at fixed prediction-list sizes of 100, 300 and 500 predicted genes. Left, mean absolute recovery for each method. Right, paired within-screen differences between AutoScreen and BioDiscovery and between AutoScreen and GPT-5.4. Points indicate mean estimates and error bars indicate 95% confidence intervals. **d,** Relative library-size reduction required to recover the same number of ground-truth hits as AutoScreen at the top-500 reference point. **e,** Overlap analysis of ground-truth hits recovered by AutoScreen, BioDiscovery and GPT-5.4 at the top 500 predictions. Bars indicate the average percentage of total hits in each intersection. **f,** Relative mean average precision at 100 predictions (MAP@100), normalized to random selection.

We compared AutoScreen in Pre-screen Design mode against BioDiscovery Agent and GPT-5.4, each given the same research goal and asked to return a ranked list of 500 genes, as well as a random baseline of 500 genes drawn uniformly from the measured gene universe. AutoScreen fused three such ranked lists produced under different anchor configurations. AutoScreen’s Pre-screen Design consistently recovered more validated targets than all baselines, with the advantage persisting across all prediction cutoffs (Fig. 2b,c). At the top 100 predictions, AutoScreen recovered 6.4% of ground-truth top 100 hits, compared with 5.4% for BioDiscovery, 3.8% for GPT-5.4, and 0.2% for 100 randomly selected genes. The margin remained at larger library sizes: at the top 300 predictions, AutoScreen recovered 10.7% of ground-truth hits compared with 9.4% for BioDiscovery and 7.0% for GPT-5.4; at the top 500 predictions, AutoScreen recovered 13.4%, compared with 12.2% and 9.1%, respectively. To recover the same number of true positives at the top-500 reference point, AutoScreen required a 50.0-fold smaller library than random selection, a 2.4-fold smaller library than GPT-5.4 and a 1.2-fold smaller library than BioDiscovery (Fig. 2d).

Overlap analysis of genes predicted by each method showed that AutoScreen recovered the largest shared and method-specific set of validated hits (Fig. 2e). The intersection among AutoScreen, BioDiscovery and GPT-5.4 top-500 hits accounted for 5.38% of total hits on average, while AutoScreen shared 3.87% with BioDiscovery and 0.86% with GPT-5.4. AutoScreen contributed a substantial unique component, recovering 3.32% of total hits not recovered by either baseline, compared with 2.03% unique to BioDiscovery and 2.02% unique to GPT-5.4. Consistent with this improved top-ranked recovery, AutoScreen achieved the highest MAP@100, with a 275.43-fold improvement over random selection, compared with 216.22-fold for BioDiscovery and 126.30-fold for GPT-5.4 (Fig. 2f). These results indicate that AutoScreen not only recovers targets shared with existing agentic and LLM baselines, but also identifies more validated targets and ranks genes more effectively near the top of its predictions.

### Knowledge-based retrieval and anchored gene emission drive target ranking

The remaining agents complete the pipeline. With items supplied by Agents 1 and 2, the Knowledge Base Search Agent (Agent 3) and Information Synthesis Agent (Agent 4) produce the ranked candidate list, and the Target Review Agent (Agent 5) annotates each candidate with cell-line-specific context to assemble the final report. The Knowledge Base Search Agent coordinates 26 specialized subagents (Table 1), routing each category of search item to the appropriate database subagent—spanning pathway, clinical, interaction, gene-knowledge and drug resources (e.g. KEGG pathway, genetic-interaction and TCGA agents)—which query via API or direct access, then parse the returned gene lists and normalize them to a common identifier space, retaining each source’s native score or evidence tag.

**Table 1.**
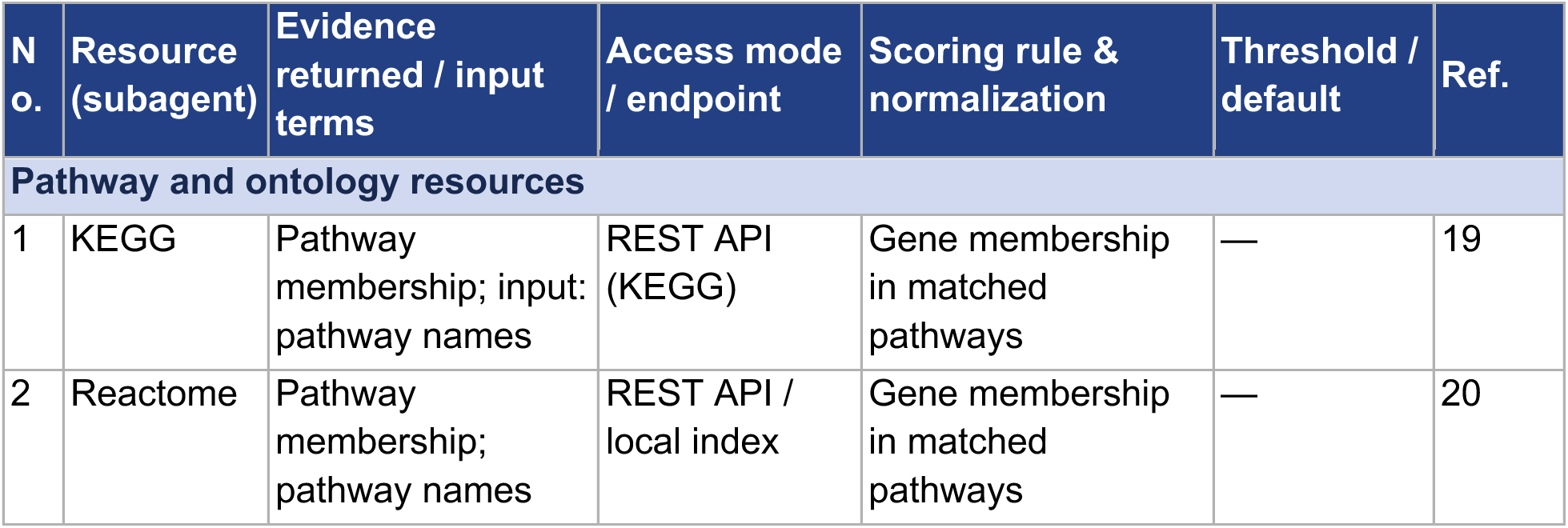
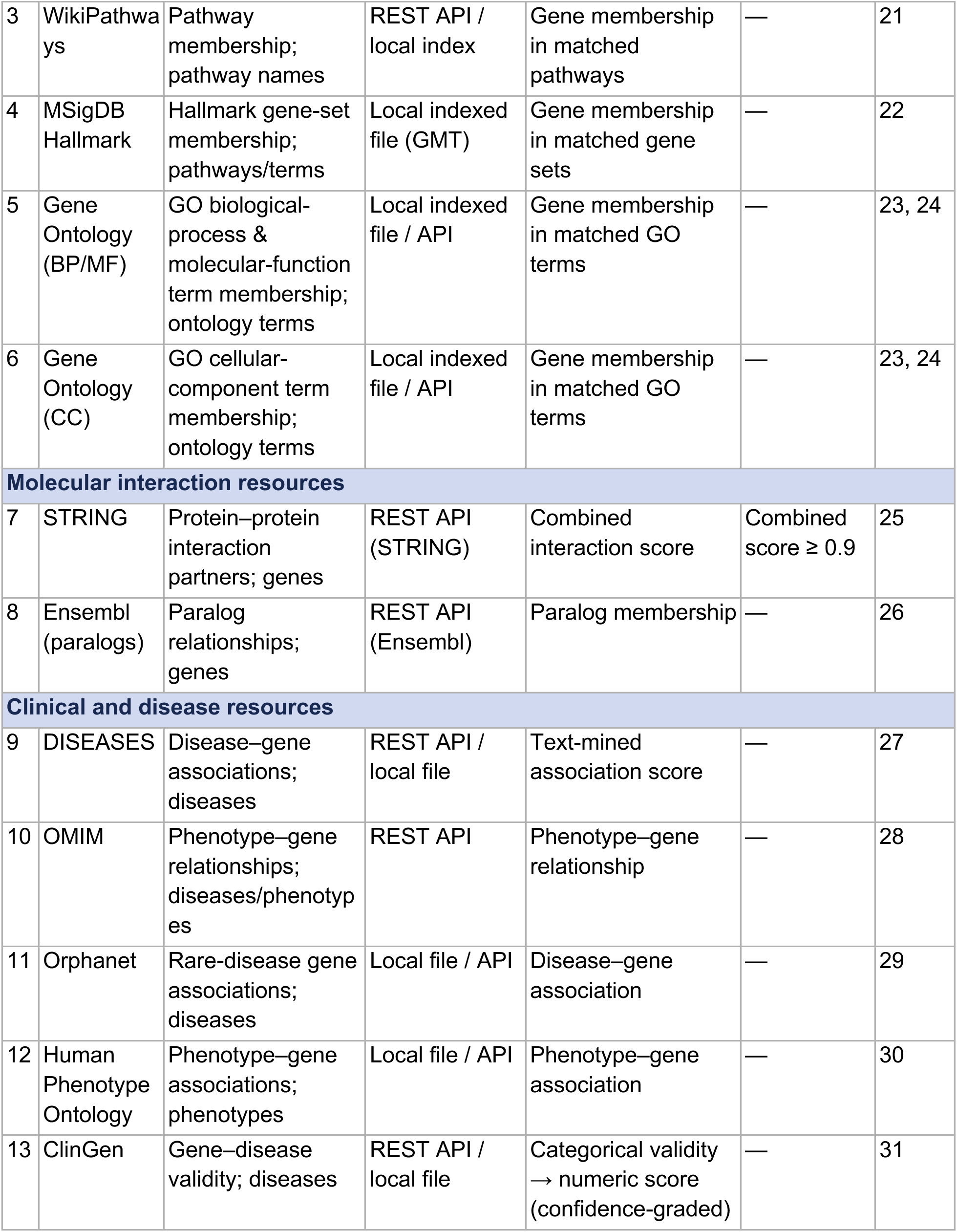
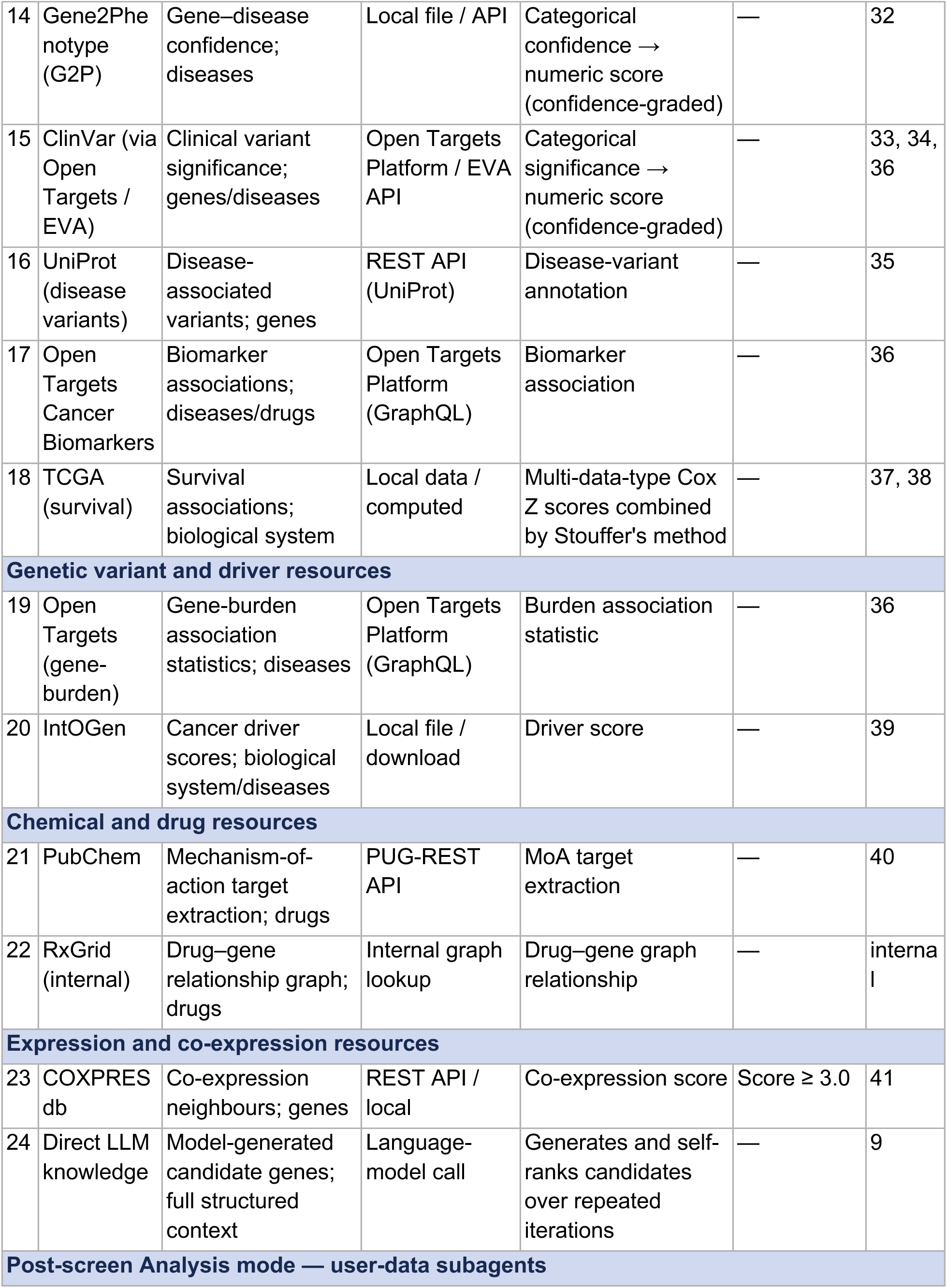
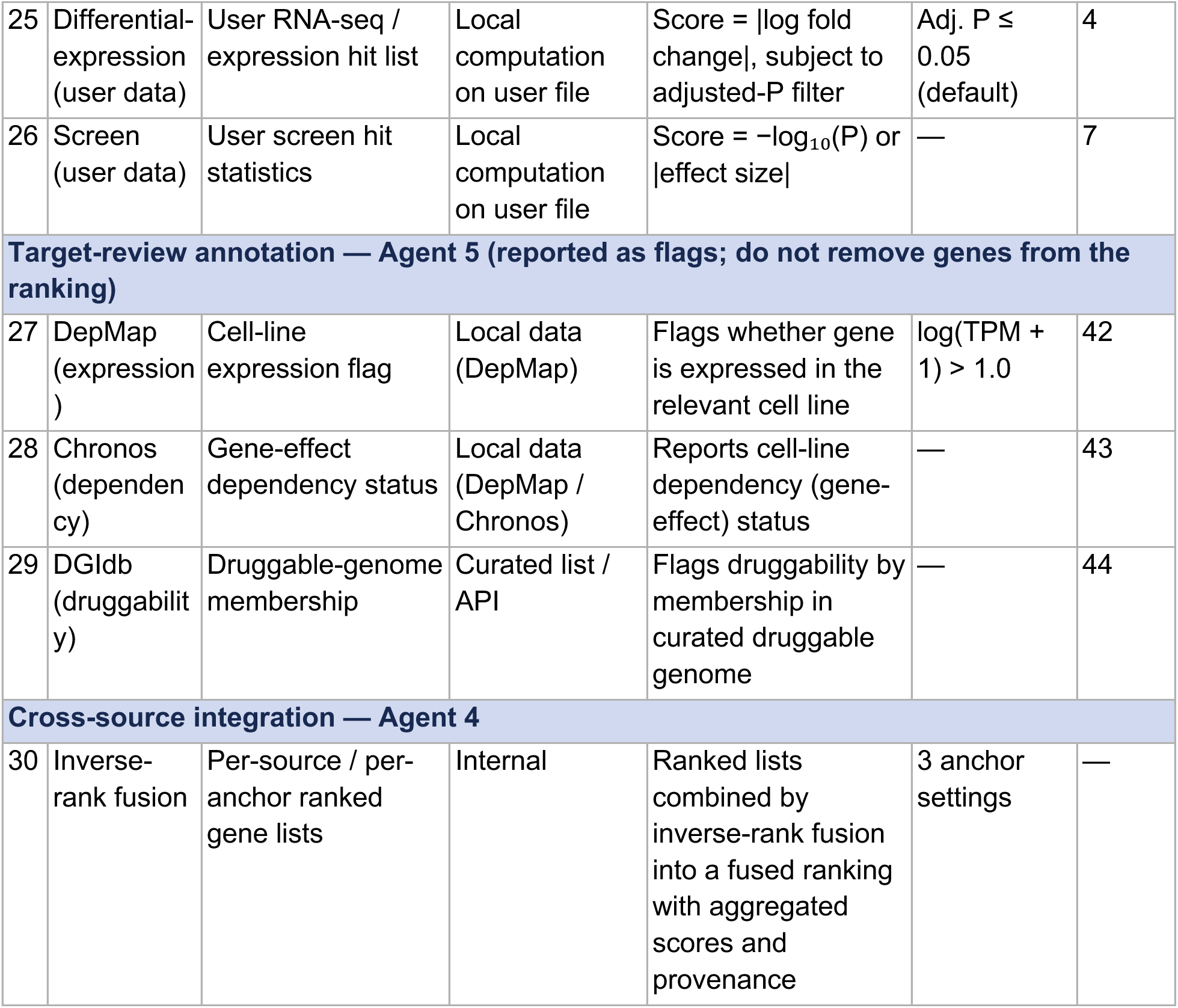
Resources used across AutoScreen agents. Part A (rows 1–26) lists the subagents of the Knowledge Base Search Agent (Agent 3), which routes each category of search term from Agent 2 to specialized subagents run in parallel — 24 knowledge-based subagents used in both modes and two additional user-data subagents activated in Post-screen Analysis mode. Continuous within-source statistics are min–max normalized across the genes returned by that source; membership-based subagents (pathway and ontology resources) contribute binary indicators; confidence-graded resources map categorical evidence levels to numeric scores before normalization. Part B (rows 27–29) lists the annotation resources reported as flags by the Target Review Agent (Agent 5); flags do not remove genes from the ranking. Part C (row 30) describes cross-source integration by the Information Synthesis Agent (Agent 4), which combines the anchor-conditioned ranked lists (n = 5, 10, 20) by inverse-rank fusion. Access modes include REST or GraphQL API, local indexed file, local computation, graph lookup, and language-model call; exact endpoint URLs and resource versions are configured per deployment. Reference numbers correspond to the manuscript reference list.

Without correction, this ‘fame bias’ depresses recovery of screen-specific candidates that sit further from the literature spotlight. The Information Synthesis Agent counteracts this bias through anchored gene emission (Fig. 1b). For each screen, the agent first synthesizes a screen-specific anti-hub anchor—a short list of textbook hub genes that the language model commonly over-predicts but that lack mechanistic relevance to the assayed phenotype—using the top-ranked categories surfaced by the knowledge bases as biological context. Critically, the anchor is derived from each screen’s knowledgebase evidence rather than being a static, predefined hub list: this couples Agent 3’s per-screen retrieval to the anchor identity, so the genes the agent is asked to avoid are themselves screen-specific rather than a universal blacklist. To enumerate candidate genes, the agent uses an anchored-emission prompt with two separable parts: a screen-specific anchor (the anti-hub avoidance list above) and a base prompt that elicits the ranked gene list. The novelty lies in the anchor and its surrounding synthesis, not in the base prompt. We insert the anchor as a soft ‘avoid’ constraint that steers predictions away from frame-biased hub genes and toward screen-specific mechanisms, and we treat the base prompt as an interchangeable component.

To test whether the improvement depended on a particular base prompt, we ran the framework with several alternatives, including one adapted from the BioDiscovery Agent. Target recovery improved in each case (Supplementary Fig. 3), indicating that the gain is driven by our framework—the anchoring and fusion steps—rather than by any single prompt template, and that the framework generalizes across different base prompts for this task. We added two additional steps around emission. To reduce sensitivity to the choice of anchor length, the agent runs the anchor-and-emission procedure independently at three anchor counts (n = 5, 10 and 20) and combines the resulting ranked lists by inverse-rank fusion (Methods). The fused list retains per-variant rank and fusion-score provenance for every emitted gene. We used comparisons against ground truth only to tune anchor design and select the reasoning backend offline, not as an online training signal. Because emission is decoupled from both the base prompt and the reasoning backend, either can be replaced without changing the retrieval or fusion pipeline.

### Ensemble fusion across hub-list configurations improves target recovery

To test whether AutoScreen’s performance depended on a single hub-list configuration, we compared candidate rankings generated from three hub lists with the ensemble ranking produced by fusing their outputs. Each individual hub list recovered validated hits, but the ensemble consistently matched or exceeded the best individual configuration across prediction-list sizes (Fig. 3a). This indicates that the improvement was not driven by a single privileged hub list, but by integrating complementary target signals across multiple configurations.

**Figure 3.**
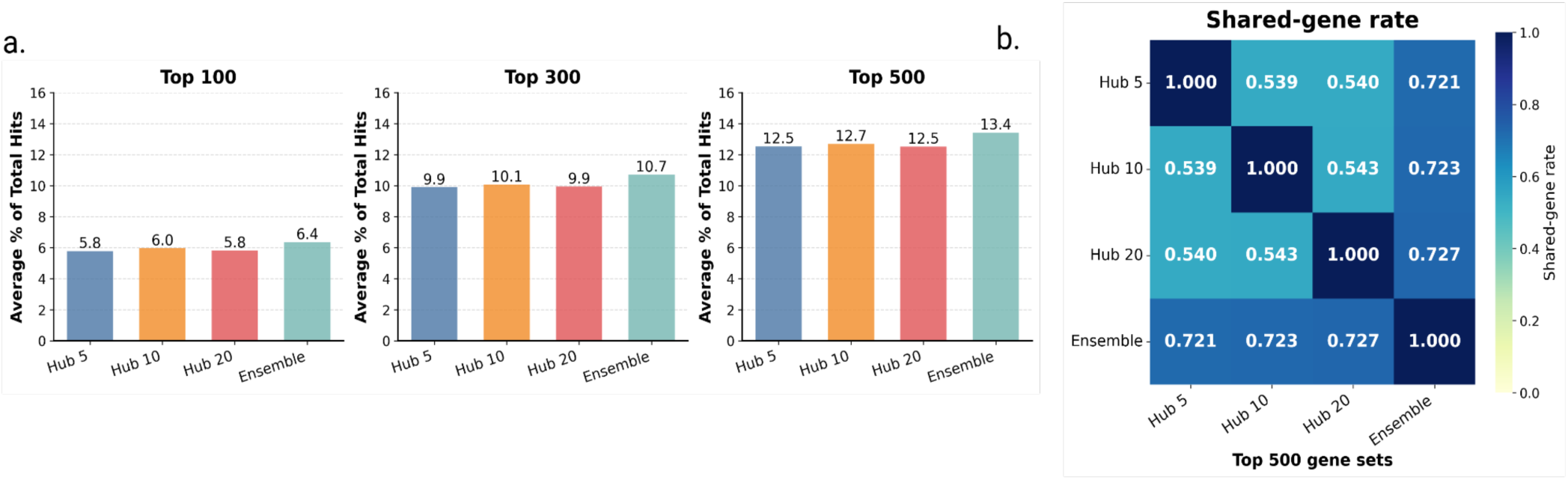
Ensemble fusion across hub-list configurations improves target recovery. **a,** Validated-hit recovery for candidate rankings generated from individual hub-list configurations and from the ensemble ranking, evaluated at the top 100, top 300 and top 500 predictions. The ensemble matched or exceeded individual hub-list configurations across prediction-list sizes. **b,** Shared-gene rate between pairs of tops-500 predicted gene sets. Individual hub-list configurations produced partially overlapping but non-identical candidate sets, whereas the ensemble retained higher overlap with each individual configuration, consistent with integration across complementary rankings.

We also quantified how much the ranked outputs overlapped. Instead of comparing summary embeddings, we measured the shared-gene rate between pairs of top-ranked results, defined as the fraction of genes in the top-K list that were common to both methods. The individual hub-list configurations shared only a moderate fraction of top-ranked genes, indicating that each produced a partially distinct candidate set (Fig. 3b). By contrast, the ensemble shared a higher fraction of genes with each individual configuration, consistent with its role as an integrated ranking that retains common high-confidence targets while incorporating candidates unique to individual hub lists. Together, these results show that varying the hub list exposes complementary target neighborhoods, and that ensemble fusion improves target recovery by combining these partially overlapping rankings. Individual hub-list configurations shared approximately half of their top-500 predictions yet showed similar mean recovery (Fig. 3a), indicating that the predictions unique to each configuration are drawn from equally productive gene neighborhoods.

### Two examples illustrate the pathway-level effect of anchored emission

In autophagy screen 1593, BioDiscovery ranked the non-hit canonical anchors MTOR and AKT1 at positions 1 and 16, whereas AutoScreen promoted canonical autophagy machinery (WIPI2, ATG2A, ATG9A, RB1CC1, ATG13) into the top 15 (Supplementary Table 1). In the K562 CTx-DTA screen (1161), BioDiscovery prioritized the ganglioside receptor pathway but did not recover the diphthamide-biosynthesis pathway required to generate DTA-targeted eEF2-diphthamide; AutoScreen ranked DPH1, DPH2, DPH5, DPH6 and DPH7 at positions 3–19 (Supplementary Table 2). These examples illustrate pathway-level rank reallocation rather than a deterministic causal effect of any individual anchor gene.

To test whether these gains depended on a particular language model, we evaluated AutoScreen across seven language-model backends—spanning both reasoning and non-reasoning models—on the 207 benchmark screens common to all configurations, comparing the Hub 5, Hub 10, Hub 20 and ensemble synthesis settings within each backend (Fig. 4). Absolute MAP@100 varied across models, reflecting differences in their underlying capability, yet within nearly every backend the ensemble setting achieved the highest MAP@100, matching or exceeding the best individual hub-anchor configuration. Because this advantage held even across non-reasoning backends, the benefit of fusing multiple context-aware synthesis settings is a property of the anchored-ensemble design rather than of any particular model. This consistency across the evaluated backends supports the robustness of the anchored-ensemble design and the treatment of the language-model component as a replaceable backend, although backend selection was performed against experimentally validated targets.

**Figure 4.**
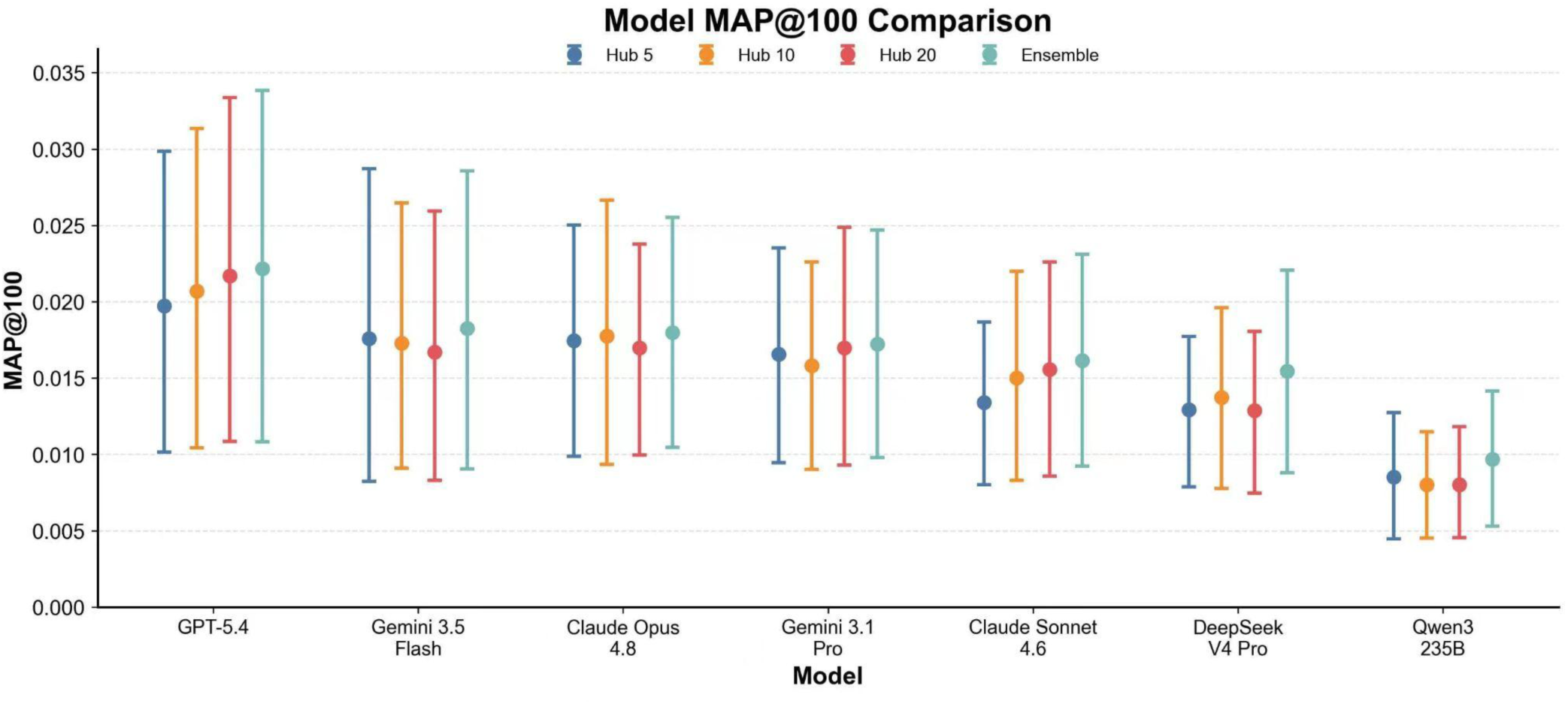
Model-dependent MAP@100 performance across AutoScreen backbones and synthesis strategies. Point-range plot comparing absolute MAP@100 across seven reasoning backends evaluated on 207 common benchmark screens. For each model, performance is shown for Hub 5, Hub 10, Hub 20, and Ensemble synthesis settings. The ensemble condition generally provides the highest MAP@100 across models, indicating that combining multiple context-aware synthesis settings improves target-ranking performance relative to individual hub-anchor settings. Points indicate mean MAP@100 across screens, and error bars indicate 95% confidence intervals estimated from 5,000 bootstrap resamples clustered by BioGRID study.

### Application and validation of AutoScreen in cancer immune evasion studies

We next applied AutoScreen to two distinct types of problem, post-screen analysis (physical experiments have been completed) vs. pre-screen design (generating hypotheses before actual experiments). These two modes are central to the scientific process of target discovery: In Post-screen Analysis mode, we used AutoScreen to re-rank experimental screen hits from two independent CRISPR activation screens probing NK-cell-mediated immune evasion in leukemia and melanoma cell lines. In Pre-screen Design mode, we used AutoScreen to construct a library directly from a study design, prioritizing candidates before any screen is run.

### Evaluating AutoScreen in post-screen analysis mode

For Post-Screen Analysis, we first performed a new surfaceome CRISPR activation screen^11,12^ in K562 cells. We transduced dCas9-SAM K562 cells with a surfaceome-protein activation CRISPRa library and co-cultured them with or without primary human NK cells, then quantified sgRNA abundance by sequencing surviving cells following NK cell-mediated killing (Fig. 5b).MAGeCK analysis identified 317 statistically significant candidates (167 enriched and 150 depleted) out of all measured genes in the library (Fig. 5c). To obtain the mechanistic context needed to prioritize follow-up, we applied Post-screen Analysis mode (Fig. 5d), with the full MAGeCK-ranked output for all measured genes and study description as input. AutoScreen’s Post-screen Analysis mode integrated the primary screening signal with literature, pathway and clinical evidence, moving MUC1, for example, from rank 118 to 5, PDPN from rank 81 to 44, and LRRC15 from rank 1384 to 659 (Fig. 5e). Individual validation using multiple gRNAs to perturb each gene confirmed that MUC1, PDPN, and LRRC15 activation all significantly increased resistance to NK-cell killing relative to non-targeting controls (Fig. 5f). These successful results indicate that AutoScreen’s re-ranking recovered genuine cancer-immune target biology that had been substantially down-ranked in the primary analysis.

**Figure 5.**
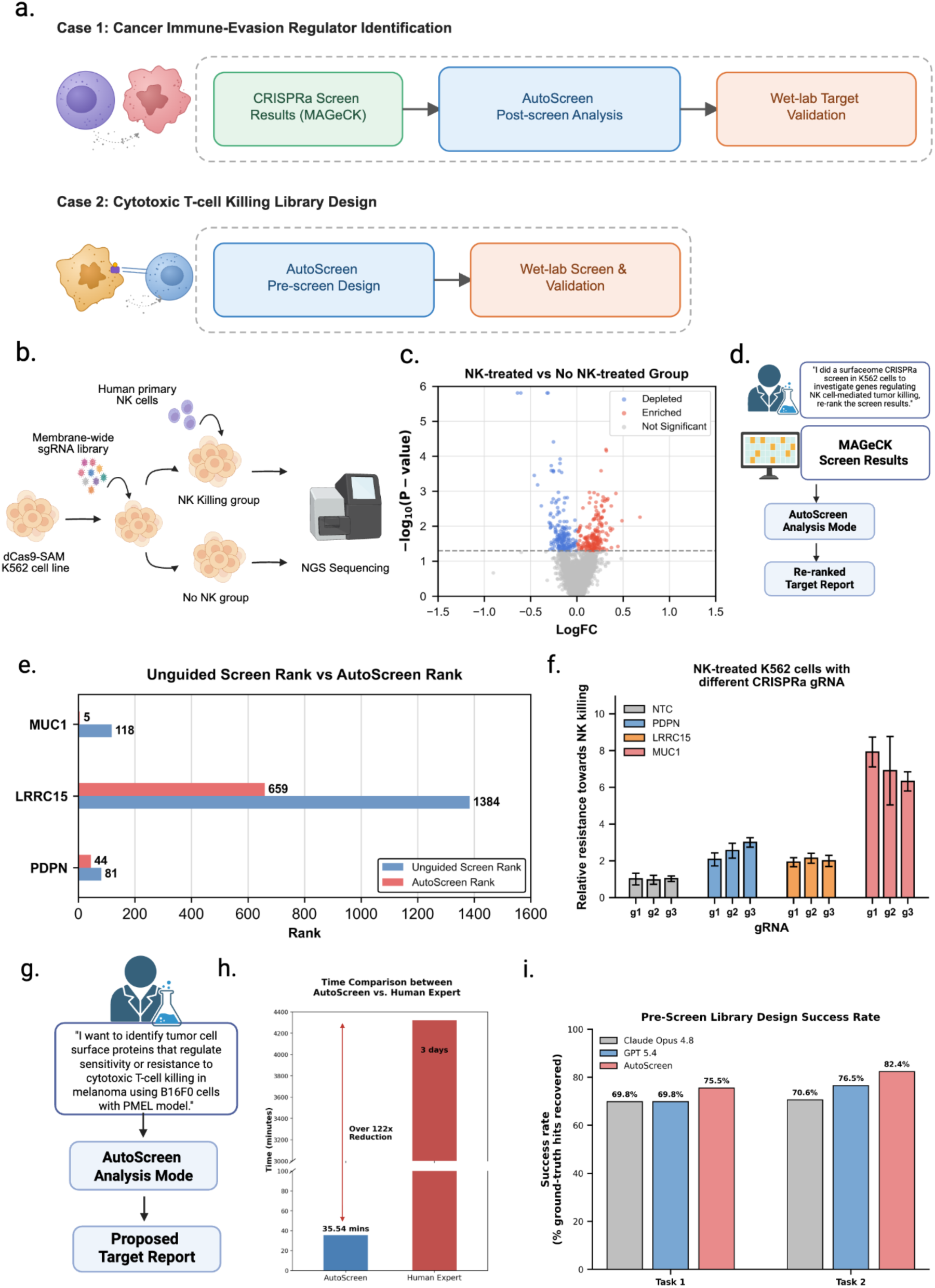
AutoScreen applications in immune evasion studies, validating the effectiveness of AutoScreen in both post-screen re-ranking and pre-screen library design. **a,** Overview of the two real-world validation cases: post-screen re-ranking of a CRISPRa screen for cancer-immunology regulator identification (top), and pre-screen library design for cytotoxic T-cell killing screens (bottom), each proceeding from screen data or study design through AutoScreen to wet-lab target validation. **b,** Schematic of the NK cell CRISPR activation screen. dCas9-SAM K562 cells transduced with a membrane activation library are co-cultured with or without primary human NK cells; genomic DNA is harvested and sequenced to quantify sgRNA abundance. **c,** Volcano plot showing differential sgRNA enrichment (log fold change vs –log₁₀ P value) between NK-treated and control conditions, analyzed with MAGeCK to identify candidate resistance (enriched) and sensitization (depleted) hits. **d,** Analysis workflow illustrating AutoScreen’s re-ranking of MAGeCK results and generation of an evidence-backed target report. **e,** Unguided screen rank (blue) versus AutoScreen re-ranked rank (red) for MUC1 (118→5), LRRC15 (1384→659), and PDPN (81→44). **f,** In vitro validation for K562 cells: Relative resistance to NK-cell killing for K562 cells expressing CRISPRa sgRNAs targeting PDPN, LRRC15 or MUC1, compared with non-targeting control (NTC) sgRNAs, across three independent sgRNAs (g1-3) per gene. Data is normalized to the NTC condition from 3 independent replicates. **g,** Design-mode workflow: AutoScreen’s Pre-screen Design mode was queried independently for each of two tumor-model/antigen-system combinations (Task1: B16F0-PMEL melanoma, Task2: MC38-OT1 colon cancer) and each returning a proposed target library. **h,** Time comparison for generating libraries: AutoScreen (automated, minutes) vs expert (manual, days). **i,** Success rate (% ground-truth hits, defined as genes independently called significant by both MAGeCK and Broad-method analysis at neg|fdr < 0.1 or pos|fdr < 0.1) of AutoScreen’s designed library compared with two general-purpose LLM baselines (Claude Opus 4.8 and GPT 5.4), across two independent CRISPR screens (Task 1 & 2).

To test whether the same re-ranking Post-screen Analysis approach generalized across an independent context, we also applied AutoScreen to a second CRISPR activation screen of dCas9-VP64 A375 melanoma cells transduced with a membrane-protein activation library and co-cultured with or without primary human NK cells (Supplementary Fig. 1a,b). MAGeCK analysis identified 186 statistically significant candidates out of all measured genes in the library, but the full ranked list lacked the mechanistic context needed to prioritize follow-up. Supplying the full MAGeCK-ranked output for all measured genes, together with the study description, to AutoScreen’s Post-screen Analysis mode (Supplementary Fig. 1c) integrated the screening signal with literature, pathway and clinical evidence and sharply re-prioritized the CEACAM family: CEACAM1 ranked highly in the original analysis (rank 2), while AutoScreen also elevated CEACAM5 (rank 143→38) and CEACAM6 (rank 155→26) despite weaker primary-screen significance, drawing on interaction-network and clinical evidence (Supplementary Fig. 1d).

### Evaluating AutoScreen in Pre-screen design mode

In Post-Screen Analysis, AutoScreen recovered more benchmark hits than general-purpose LLM baselines while generating candidate lists substantially faster than manual expert curation. To test whether the same system could design a candidate library *de novo*, we queried AutoScreen’s Pre-screen Design mode independently in two test studies. First, in task 1, AutoScreen generated hypotheses for a target discovery library in the context of using melanoma cells (B16F0) to screen against cytotoxic T-cells bearing PMEL T-cell-receptor. Second, in task 2, similar tasks were performed for a target screen using colon adenocarcinoma cells (MC38) against the OT-I T-cell system. In both cases, we did not supply any prior screening data (Fig. 5g).

To generate ground truth for prospective benchmarking, we performed two new, unpublished CRISPR screens for these settings in the wet lab. AutoScreen generated each of these libraries in 35.54 minutes on average, compared with approximately 3 days for a human expert performing the curation manually, an over 122-fold reduction in time (Fig.5h). Each task-specific query returned a proposed candidate library, which was then benchmarked against the corresponding CRISPR screen’s measured hits. Both screens used a shared focused CRISPR library of 3,340 genes, and each method was asked to nominate up to 500 candidate genes. Ground truth was defined as genes independently called significant by both MAGeCK and Broad GPP (Genetic Perturbation Platform) gene scoring analysis (neg|fdr < 0.1 or pos|fdr < 0.1), yielding 53 ground-truth hits for B16F0-PMEL and 17 for MC38-OT1. Across both tasks, AutoScreen’s designed libraries recovered more ground-truth hits within their predictions than either general-purpose LLM baseline: 75.5% versus 69.8% (Claude Opus 4.8) and 69.8% (GPT 5.4) in the B16F0-PMEL screen, and 82.4% versus 70.6% and 76.5% in the MC38-OT1 screen (Fig. 5i).

### The AutoScreen Resource Hub provides pre-computed analyses for the community

To scale AutoScreen across large functional-genomics analyses, we built the AutoScreen Resource Hub as a reusable evidence layer indexed by canonical knowledgebase identifiers (Fig. 6). We populated the hub by running AutoScreen in Pre-screen Design mode over more than 500 public functional-genomics datasets spanning CRISPR screens from BioGRID ORCS, transcriptomic studies from Expression Atlas and GWAS traits from the UK Biobank. For each new task, the pipeline structures the user query and optional data into screen-relevant terms — genes, diseases or pathways, GO and phenotype terms, and drugs — then matches them to canonical knowledge-based identifiers such as Reactome, GO, EFO or MONDO IDs, which define the cache keys used for downstream retrieval; the full hit/miss logic is described in Fig. 6. The hub integrates 24 knowledge sources spanning pathway, interaction and disease annotations, providing a median of 82 curated entries from 18 sources per screen (Supplementary Fig. 4).

**Figure 6.**
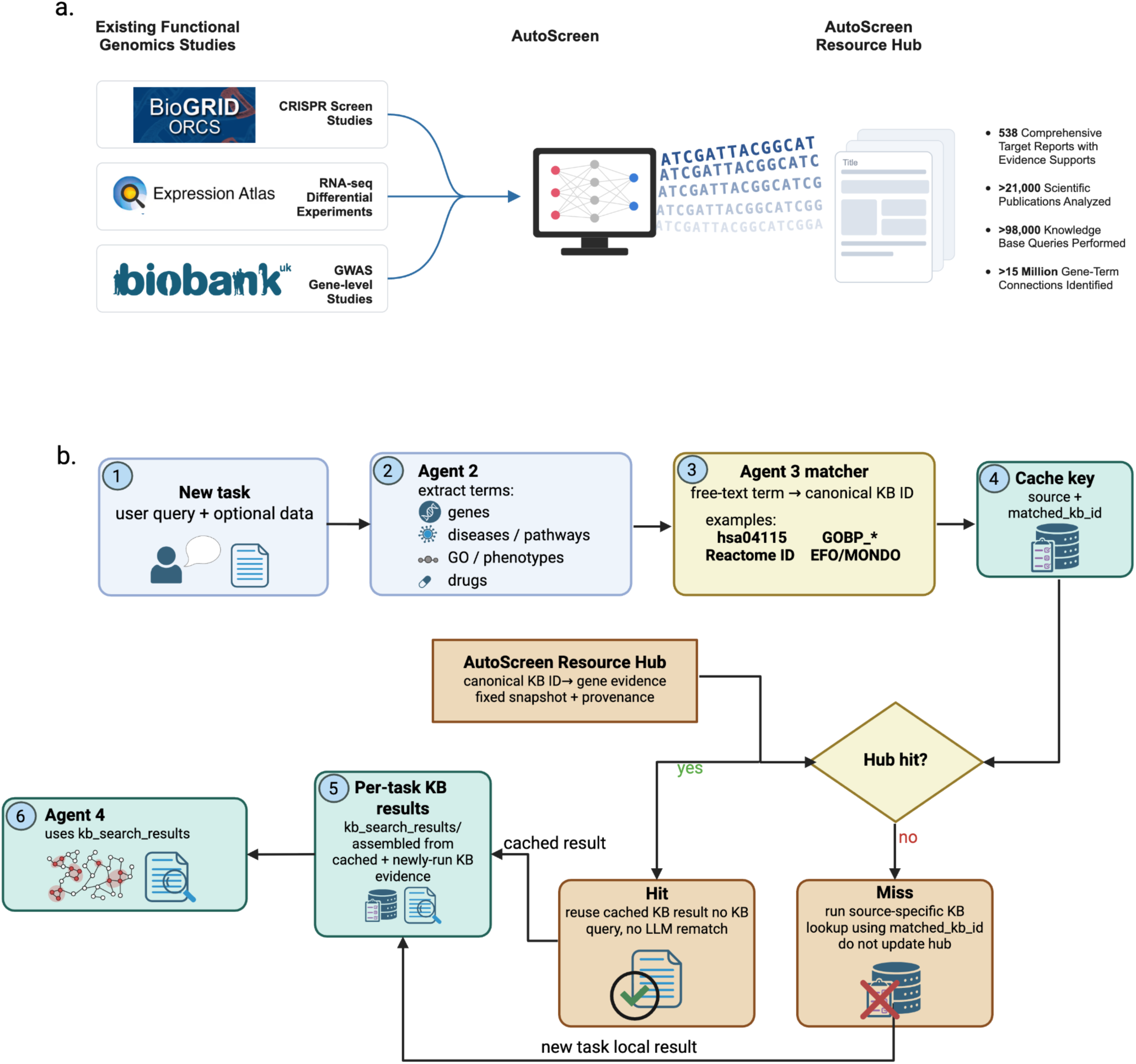
Building and using the AutoScreen Resource Hub for scalable functional genomics analysis and as an evolving, reusable benchmark. **a**, Overview of the AutoScreen Resource Hub: public CRISPR (BioGRID ORCS), transcriptomic (Expression Atlas) and GWAS (UK Biobank) datasets are processed by AutoScreen into a cached evidence layer, comprising 538 comprehensive target reports with evidence support, drawn from over 21,000 scientific publications, more than 98,000 knowledge-base queries and over 15 million gene-term connections. b, Workflow for cached knowledge synthesis across large-scale benchmark analyses. A user query and optional data are first structured by Agent 2 into genes, diseases/pathways, GO/phenotype terms and drugs. These free-text terms are then matched to canonical knowledge-based identifiers — such as KEGG pathway IDs, GO term accessions, Reactome IDs and EFO/MONDO disease codes — by a term-matching step within Agent 3, forming a cache key of source and matched identifier. Matched identifiers are checked against the AutoScreen Resource Hub, which stores fixed, provenance-linked gene evidence indexed by canonical KB identifiers. Cache hits reuse existing knowledge-based results without repeated source-specific lookup or LLM rematching, whereas if cache misses, this would trigger a source-specific knowledge-based search; missed results are still retained and kept as task-local results (and not written back to the hub). Cached hits and newly generated local results are then assembled into per-task knowledge/results, and passed to the Agent 4 synthesis step to produce context-aware candidate-gene rankings. This entire process can be run iteratively, continuously, as new resources become available.

The Resource Hub stores fixed, provenance-linked gene evidence for matched canonical identifiers. Cache hits reuse existing knowledgebase results without repeated source-specific lookup or LLM rematching; cache misses trigger a source-specific knowledgebase search, and miss results are kept as task-local results that are not written back to the hub. Cached and newly generated results are then assembled and passed to the information synthesis step, which adapts the evidence to the biological context of each task to produce candidate-gene rankings. This design separates stable, reusable evidence retrieval from task-specific synthesis, reducing redundant knowledge-base queries while preserving transparent evidence provenance for each ranked candidate list. Further, this efficient AutoScreen inference approach helps to set up this Resource Hub as a continuously evolving, updating, and expanding benchmark.

## Discussion

AutoScreen addresses the prioritization bottleneck in functional genomics by integrating language-model reasoning, multi-source knowledge retrieval and context-aware anchored gene emission within a modular multi-agent framework. Across 320 genome-scale CRISPR screens it recovered more validated hits at smaller library sizes than statistical scoring, general-purpose LLM and agent baselines. In the immune-evasion case studies using newly generated wet lab screen datasets, AutoScreen helped to prioritize experimentally confirmed target genes, and prospectively designed libraries that recovered more screen hits vs. alternative baseline methods. These results indicate that synthesizing multi-modal evidence, rather than relying on any single source, is what drives AutoScreen’s target discovery performance—an interpretation supported by the consistent margin of improvement, when comparing the integrated approach vs. literature, web search, and LLM sources.

The system’s two modes map onto distinct stages of the discovery pipeline. Post-screen Analysis re-weights existing screen data against external evidence: in a leukemia surfaceome screen, re-ranking recovered experimentally validated NK-resistance hits (MUC1, PDPN, LRRC15) that had been lower-ranked by the statistical method, and in an independent melanoma screen it elevated the CEACAM family (CEACAM1, CEACAM5 and CEACAM6), which is currently under clinical investigation. Pre-screen Design constructs candidate libraries directly from a study description, without access to any prior screening data. Across two independent cytotoxic-T-cell-killing screens used for prospective validation, AutoScreen’s designed libraries recovered more hits than either general-purpose LLM baseline in both screens: 75.5% versus 69.8% and 69.8% in one screen, and 82.4% versus 70.6% and 76.5% in the other, while requiring a small fraction of the time needed for equivalent manual curation. Together these modes make rigorous target selection less dependent on individual domain expertise and curation time.

A defining feature of AutoScreen is that its recommendations are auditable at multiple levels. The pipeline records the anchor variant and fusion step that produced each ranking, and preserves the knowledgebase evidence—spanning pathway, interaction and disease sources—consulted during analysis (Supplementary Fig. 4). As illustrated by the pathway-level rank reallocation in screens 1593 and 1161 (Supplementary Tables 1, 2), this design makes it possible to trace how anchored emission shifted predictions away from frame-biased hub genes and toward screen-relevant biology. We regard this auditability, rather than predictive accuracy alone, as a prerequisite for routine use of AI systems in experimental design.

The modular separation of retrieval and synthesis carries a broader methodological implication. Target prioritization should be evaluated as a task in its own right, rather than inferred from general-purpose reasoning benchmarks, since improvements on general leaderboards do not straightforwardly translate to candidate selection in specific biological contexts. Because the synthesis agent accepts interchangeable language-model backends, new model families can be benchmarked offline against experimentally validated targets and substituted without altering the retrieval or fusion pipeline as language models advance.

Several limitations remain. AutoScreen depends on existing structured knowledgebases, which are biased toward well-studied genes and pathways and may under-represent poorly annotated biology; its outputs reflect these gaps. Thus, a future goal is to have a tunable system to help with addressing the diverse need of the system on finding “plausible” vs. “completely novel” hits. Also, our benchmark defines ground truth as the top-ranked genes from each original analysis, which rewards agreement with prior experimental data calls and may undervalue completely novel candidates. The ∼25% of AutoScreen hits unique to the system were not independently validated at scale yet, and the wet-lab validations are still small-scale tests, though successful.

Extending AutoScreen will require both broader evidence and deeper reasoning. Incorporating modalities beyond CRISPR, RNA-seq and GWAS would broaden biological coverage, while support for combinatorial designs would address phenotypes that require coordinated perturbation of multiple genes. Recursive, multilayer search—in which prioritized targets seed further exploration of connected entities—could surface higher-order relationships. Evaluation criteria that weight novelty and mechanistic diversity alongside recall would let researchers tune selection toward unexplored biology. As biological knowledgebases and reasoning models continue to grow, a transparent and modular framework of this kind offers a route to keeping target discovery aligned with the expanding literature.

## Methods

### AutoScreen system overview

AutoScreen is a modular multi-agent system that transforms a free-text research description into a ranked, evidence-backed list of candidate genes with pipeline provenance. The system is organized as five specialist agents that execute in sequence—a Deep Research Agent (Agent 1), an Information Restructure Agent (Agent 2), a Knowledge Base Search Agent (Agent 3), an Information Synthesis Agent (Agent 4) and a Target Review Agent (Agent 5)—each consuming the structured output of the preceding stage. The pipeline runs in two modes. In Pre-screen Design mode, AutoScreen receives only a study description and constructs a *de novo* candidate library. In Post-screen Analysis mode, it additionally ingests an experimental hit list and the associated statistical metrics (for example, *P* values and log fold changes from MAGeCK or DESeq2) and re-ranks the measured genes by integrating the screening signal with external evidence. Each agent communicates through structured JSON, and every gene in the final report is accompanied by the retrieval records and the ranking-pipeline trace that surfaced it. Agents are coordinated by a central orchestrator (run_pipeline) that supports both single-query and batch execution and writes per-stage outputs to disk for auditability.

The system is implemented in Python. Language-model calls are issued through a unified interface that maps a backend identifier to a provider, model snapshot and decoding configuration, allowing any OpenAI-compatible chat model to be substituted without changes to agent logic. Unless otherwise stated, Agents 1-5 used GPT-5.4 as the reasoning backend; backend selection was performed offline against experimentally validated targets (see “Reasoning backends” below). API requests were distributed across a rotating pool of keys to support large-scale batch evaluation.

### Deep Research Agent (Agent 1)

The Deep Research Agent transforms the user description into a literature-grounded summary. It first refines the raw query into a search-engine-compatible query with a single deterministic language-model call (temperature 0, JSON output), then retrieves evidence in parallel from multiple providers—academic search (Google Scholar via SerpAPI), a web research provider (Perplexity) and the language model’s own parametric knowledge. Each provider’s raw output is summarized independently and then integrated into a single narrative through a further model call. A final review pass validates each sentence of the integrated summary against the retrieved sources and emits a critique together with a revised answer. The agent records every source with its title, URL, authors and publication date, and reports the total number of articles consulted; these source records seed the provenance chain carried through the rest of the pipeline.

### Information Restructure Agent (Agent 2)

The Information Restructure Agent distills the narrative summary and original query into ontology-aligned search terms. Using a reasoning model (GPT-5.4) with JSON output, it decomposes the evidence into eight controlled categories: biological system (one cell-line, cell-type, tissue or organ name standardized from the input text), cellular components, genes (validated HGNC symbols)^18^, pathways (Reactome/KEGG names), ontology terms (Gene Ontology), diseases, phenotypes and drugs. Extracted entities are normalized to canonical identifiers, and invalid or unsupported terms are removed. The agent then performs a configurable number of critique rounds (two by default), in which each round re-validates and corrects the previous structured output against the source text, removing unsupported entries and adding missing entities. The output is a structured term set that routes the downstream database search.

### Knowledge Base Search Agent (Agent 3)

The Knowledge Base Search Agent retrieves candidate genes from 26 specialized resources in parallel. Each category of search term produced by Agent 2 is dispatched to the appropriate subagents, which are executed concurrently with a thread pool; subagents with no applicable input terms are skipped. Each subagent queries its resource (by REST API, local indexed file, graph lookup or, where appropriate, a language-model call to match free-text terms to controlled vocabulary), parses the returned records into gene-level entries, and converts source-specific statistics into a within-source relevance score that is min–max normalized across the genes returned by that source. The agent writes one JSON file per source, each mapping genes to their evidence entries, scores and metadata.

The 26 subagents span six categories of evidence. Pathway and ontology resources comprise KEGG, Reactome, WikiPathways, MSigDB Hallmark gene sets, Gene Ontology biological-process/molecular-function terms and Gene Ontology cellular components.19-24 Molecular interaction resources comprise STRING protein–protein interactions (combined-score threshold ≥ 0.9) and Ensembl paralog relationships.25,26 Clinical and disease resources comprise the DISEASES database, OMIM, Orphanet, the Human Phenotype Ontology, ClinGen gene–disease validity, Gene2Phenotype, ClinVar (via Open Targets/EVA), UniProt disease variants, Open Targets Cancer Biomarkers and TCGA survival associations (multi-data-type Cox Z scores combined by Stouffer’s method).27-38 Genetic variant and driver resources comprise Open Targets gene-burden association statistics and IntOGen cancer driver scores.36,39 Chemical and drug resources comprise PubChem mechanism-of-action target extraction and an internal drug–gene relationship graph (RxGrid).40 Expression and co-expression resources comprise COXPRESdb co-expression41 (score threshold ≥ 3.0) and a direct language-model knowledge subagent that generates and self-ranks candidate genes over repeated iterations. In Post-screen Analysis mode, two optional user-data subagents are activated only when the corresponding input table is supplied. The differential-expression subagent accepts a precomputed gene-level differential-expression table, for example from bulk RNA-seq, filters genes by adjusted P value (default <0.05), and converts the supplied absolute-effect values into normalized within-source scores. CRISPR-screen results are handled separately by the screen-statistics subagent, which scores genes using −log₁₀(P) or absolute effect size, depending on the fields supplied. Confidence-graded resources (for example ClinGen, Gene2Phenotype and ClinVar) map categorical evidence levels to numeric scores before normalization. A complete list of resources, endpoints and scoring rules is provided in Table 1.

### Information Synthesis Agent (Agent 4): anchored gene emission and ensemble fusion

Agent 4 converts Agent 3’s per-source evidence into screen-specific biological context, synthesizes an anti-hub anchor list, and emits ranked candidate genes through iterative prompting. Three anchor sizes (n = 5, 10, 20) produce independent emission variants, which are combined by reciprocal-rank fusion into a single ranked list. In Post-screen Analysis mode, an additional reranking step integrates experimental screen statistics. Each stage is described below.

### Knowledge-based context

For each screen, the agent first builds a compact biological context from Agent 3’s output by ranking each source’s categories by entry count and retaining the top categories per source (three by default; categories with fewer than two entries are dropped, and the context is capped in length to fit the downstream prompt). This context summarizes the screen-specific biology surfaced by the knowledge bases.

### Anchor synthesis

Conditioned on the screen description and this knowledge-based context, a single language-model call proposes the anchor: a list of *n* textbook hub genes that language models commonly over-predict but that are not mechanistic for the specific screen. Because the anchor is derived from each screen’s retrieved evidence rather than from a fixed, predefined hub list, the genes the model is asked to avoid are themselves screen-specific. If the call fails or returns too few valid symbols, a fixed fallback anchor is used.

### Post-screen re-rank/re-prioritization component

In Post-screen Analysis mode, the agent additionally re-ranks the measured candidate pool against the anchor-conditioned priors. This step consumes three inputs: the experimental screen statistics, the study description, and the contextual priors from the preceding synthesis stage. The screen statistics define a MAGeCK-ranked candidate pool, while the priors are passed as two auxiliary context blocks — a hub/anchor caution block and a variant-specific recommendation block. Operating within this fixed pool, the agent re-ranks the measured candidates and aggregates the variant-specific rerankings by inverse-rank fusion into a final ensemble ranking. This step is inactive in Pre-screen Design mode.

### Target Review Agent (Agent 5)

The Target Review Agent annotates each candidate with cell-line-specific context to assemble the final report. Using DepMap data^42^, it flags whether each gene is expressed in the relevant cell line (log₁(TPM + 1) > 1.0) and reports its dependency (Chronos gene-effect)^43^ status, these annotations are available only for established cell lines with an exact DepMap match, and are reported as unavailable for primary cells or unmatched models. It marks druggability by membership in a curated druggable-genome list (DGIdb).^44^ These annotations are reported alongside each candidate but do not remove genes from the ranking. Optionally, the agent performs an automated literature review for the top-ranked candidates (10 by default) using a deep-research model with web search, returning for each gene a support call, a short evidence summary and citations; reviews are parallelized with per-gene time limits and cached. The final report is a table in which each gene carries its number of supporting sources, the source names, its fused relevance score, the expression/essentiality/druggability flags and, where computed, its literature support and citations.

### Reasoning backends

AutoScreen treats the reasoning component as a replaceable module rather than a trained model. The unified language-model interface maps a backend identifier to a provider and model snapshot, and the synthesis and review agents call the model directly, so new model families can be substituted without changes to retrieval or fusion. Backend selection was performed offline by comparing candidate rankings against experimentally validated targets on the benchmark described below; this offline comparison was used only for backend selection and anchor design and did not constitute an online training signal.

### Benchmark construction

We assembled a large-scale benchmark of genome-scale CRISPR screens from BioGRID ORCS. Screens were retained if they met stringent inclusion criteria (complete metadata, a condition- or phenotype-specific contrast, and a complete ranked gene result) spanning loss-of-function (CRISPRko, CRISPRi) and gain-of-function (CRISPRa) modalities and addressing drug resistance, chemical response and viral infection. We got 320 retained screens. For each retained screen we defined the top 100 genes from the original screen analysis as the ground-truth hit set, and we converted the screen metadata into a user-style, free-text target-discovery query (for example, “*study how 2-day interleukin-1β exposure affects A375 tumor resistance to NK killing*”). Each method received the same query per screen and returned a ranked gene list restricted to the screen’s measured genes.

### Baselines

AutoScreen was compared against three baselines, each given the identical per-screen query. The BioDiscovery Agent baseline used ranked gene lists produced by that system. The general-purpose LLM baseline queried a single language model in a zero-shot setting (temperature 0, JSON output), requesting a ranked list of the desired length with continuation calls when the response was truncated. The random baseline sampled gene symbols uniformly from HGNC-approved protein-coding genes with a fixed seed. All baselines emitted ranked lists in the same format as AutoScreen and were evaluated identically.

### Evaluation metrics

For each screen and each method we computed recovery as a function of prediction-list size *K*. At a given *K*, recall was the fraction of ground-truth hits contained in the top-*K* predictions and precision was the fraction of the top-*K* predictions that were ground-truth hits; curves were evaluated at *K* in steps of 10 up to 100 and steps of 50 thereafter, and summarized at *K* = 100, 300 and 500. Ranking quality near the top of the list was quantified by mean average precision at 100 predictions (MAP@100) and reported relative to the random baseline (each method’s MAP divided by the random MAP); we also computed normalized discounted cumulative gain and mean reciprocal rank. Library-size efficiency was quantified as the relative reduction in library size required to recover the same number of ground-truth hits as AutoScreen at the top-500 reference point, obtained by interpolating each baseline’s recovery curve. Method overlap was assessed at the top 500 predictions by computing, per screen, the ground-truth hits recovered by each method and the genes shared across method intersections and unique to each method, averaged across screens. All per-screen metrics were aggregated across screens as means.

### Plasmids and library construction

Plasmids for dCas9-SAM system (pLV-EF1a-dCas9VP64_2A_EGFP_2A_BSD and pLV-EFS-PCP_p65_HSF1_2A_mcherry_HygR) and library backbone vector pLV-U6_sgRNA_pp7-EFS-Puro_2A_BFP were constructed as previously described.^46^ For the human surfaceome library, eight sgRNAs per gene targeting each human cancer surface protein gene were designed using the CRISPick tool (Broad Institute; https://portals.broadinstitute.org/gppx/crispick/public). For the mouse surfaceome library, human genes were first converted to their mouse homologs using the babelgene R package, and eight sgRNAs per gene were subsequently designed using CRISPick. Oligonucleotide pools were synthesized with flanking Esp3I/BsmBI restriction sites and PCR amplification handles (Genscript) and cloned into the backbone vector pLV-U6_sgRNA_pp7-EFS-Puro_2A_BFP by Golden Gate assembly.

### Cell culture, stable cell line generation and primary cell isolation

293FT cells ( #R7007) were purchased from Thermo Fisher Scientific. B16F0 cells (#CRL-6322) and A375 cells (ATCC CRL-1619) were purchased from ATCC. K562 cell line (ATCC CRL-243) was a gift from Michael Cleary (Stanford University). MC38 cell line was a gift from John Sunwoo (Stanford University). 293FT, B16F0, A375 and MC38 cells were grown in DMEM supplemented with 1% Pen-Strep, 1% NEAA and 10% FBS at 37°C. K562 cells were grown in RPMI supplemented with 1% Pen-Strep, 1% NEAA, and 10% FBS at 37°C.

Lentiviruses were generated by transfecting 293FT cells at 80% confluence using JetOptimus and a DNA ratio of 2:2:1 transfer plasmid:pCMV-dR8.91:pCMV-VSVG according to manufacturer’s protocol. K562, A375, B16F0 and MC38 CRISPRa lines were generated by first transducing cells with pLV-EF1a-dCas9VP64_2A_EGFP_BSD and selecting with blasticidin for 3-7 days. The cells were then transduced with pLV-EF1a-PCPp65HSF1_2A_mcherry_HygR and selected in hygromycin.Cells with highly stable GFP and mCherry expression that also maintained high levels of CRISPR activation were selected for screening.

PBMCs were isolated from LRS chambers by Ficoll-Paque density-gradient centrifugation. Primary NK cells were enriched using the EasySep NK Cell Isolation Kit (StemCell) and cultured in CTS NK-Xpander medium supplemented with 5% Immune Cell Serum Replacement, 1,000 IU/mL IL-2 and 20 ng/mL IL-15. CD8⁺ T cells were isolated from the spleens of OT-I or pmel-1 transgenic mice by negative-selection magnetic enrichment and cultured in RPMI 1640 supplemented with 10% FBS, 1% Pen-Strep, 2 mM L-glutamine and 50 μM β-mercaptoethanol. T cells were activated with anti-CD3/CD28 antibodies and expanded with recombinant mouse IL-2 cytokine.

### CRISPR screens

CRISPRa K562 and A375 cells were transduced with human, and B16F0 and MC38 cells with mouse, surface protein library lentiviruses at an MOI of 0.3. After 48 h, cells were selected with puromycin for 7–10 days until ≥98% BFP⁺, as determined by flow cytometry.

For NK cell killing screens, K562 and A375 library cells were cultured with or without primary NK cells at 1,000× library coverage. For T cell killing screens, B16F0 and MC38 library cells were similarly cultured with or without mouse T cells. After 24 h, surviving cancer cells were collected and FACS-sorted for GFP⁺/mCherry⁺ cells. Genomic DNA was extracted for NGS, and screen hits were ranked using MAGeCK-RRA and Broad GPP STARS.

### Cytotoxicity assay

Leukemia K562 cells bearing CRISPRa epigenetic gene-editing system were transduced with individual gRNAs targeting PDPN, LRRC15, or MUC1 were seeded at 1 × 10⁵ cells per well. Primary NK cells were added at an E:T ratio of 2:1, with parallel wells without NK cells as controls. Cell viability was monitored every 2 h using an IncuCyte live-cell analysis system, and fluorescence measurements at 24 h were used to compare cytotoxicity among groups.

