## Supplementary Information for "AutoScreen: AI Co-Scientist System for Target Discovery in Functional Genomics"

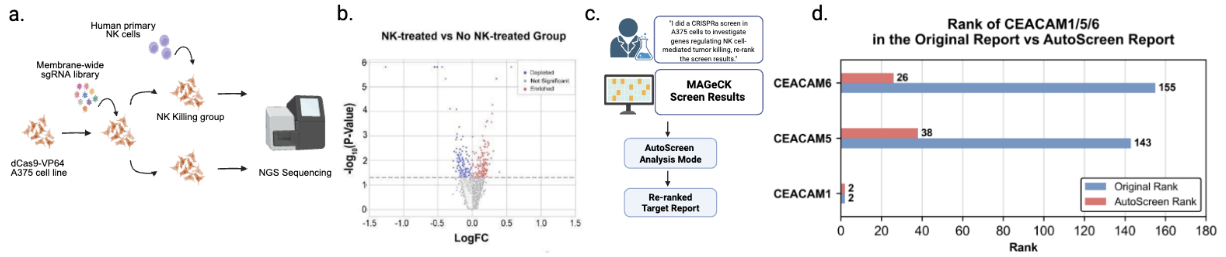


**Supplementary Figure 1 | Immune evasion in melanoma (Post-screen Analysis). a,** Schematic of the NK-cell CRISPR activation screen. dCas9-VP64 A375 melanoma cells transduced with a membrane-wide sgRNA library are co-cultured with or without primary human NK cells. Genomic DNA is harvested and sequenced (NGS) to quantify sgRNA abundance in the NK-killing and no-NK control groups. **b,** Volcano plot showing differential sgRNA enrichment between NK-treated and non-NK- treated A375 cells, analyzed with MAGeCK to identify candidate resistance (enriched) and sensitization (depleted) hits. **c,** Analysis workflow: the study goal ("I did a CRISPRa screen in A375 cells to investigate genes regulating NK cell-mediated tumor killing, re-rank the screen results.") together with the MAGeCK screen results are supplied to AutoScreen's Post-screen Analysis mode which returned as re-ranked target report. **d,** Rank of CEACAM1, CEACAM5, and CEACAM6 in the original MAGeCK-based report (blue) and the AutoScreen re-ranked report (red): CEACAM1 (2→2), CEACAM5 (143→38), and CEACAM6 (155→26).

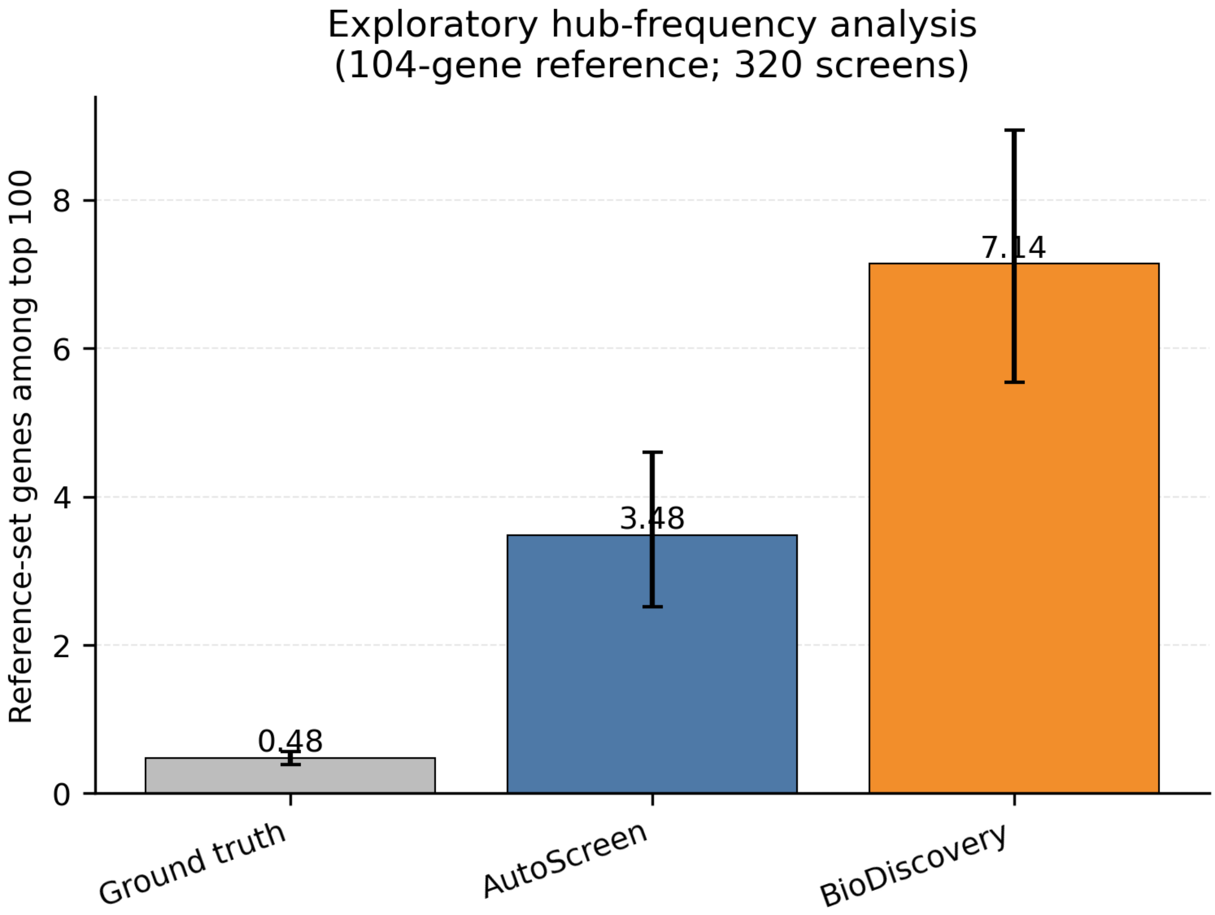


**Supplementary Figure 2 | Hub-gene frequency analysis.** Mean number of reference-set genes among the top 100 predictions for ground truth, AutoScreen and BioDiscovery Agent across 320 benchmark screens, with 95% study-cluster bootstrap confidence intervals. The reference set comprises 104 genes selected by hub score = $freq\_KB(g) \times (1 - freq\_hit|KB(g))$, computed across 25 knowledge-based sources and 320 screens, capturing genes that are ubiquitous in structured databases but rarely validated as screen-specific hits.

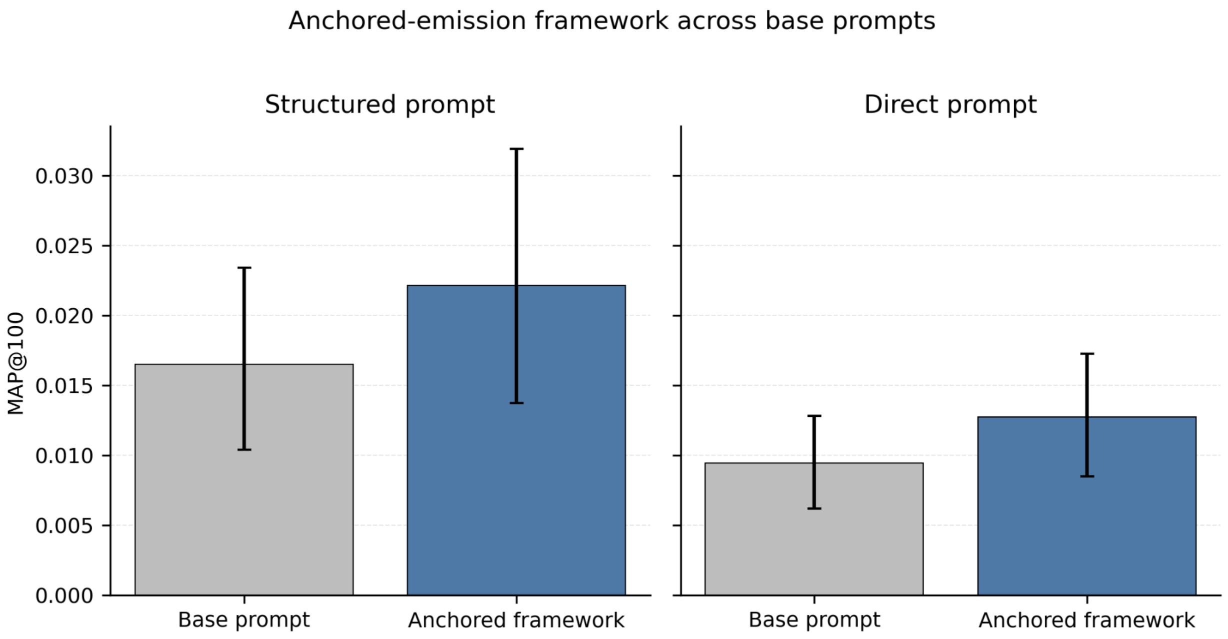


**Supplementary Figure 3 | Anchored-emission framework performance across base prompts.** Each panel compares a base prompt used as a single emission with the same prompt embedded in the anchored-emission framework, which generates three rankings using screen-specific anchors of 5, 10 and 20 genes and combines them by inverse-rank fusion. Left, a structured prompt with iterative Reflection–Plan–Solution steps adapted from the BioDiscovery Agent; right, a direct prompt requesting a ranked gene list without multi-step reasoning. Both panels use the same 320 screens evaluated on top-100 ground truth.

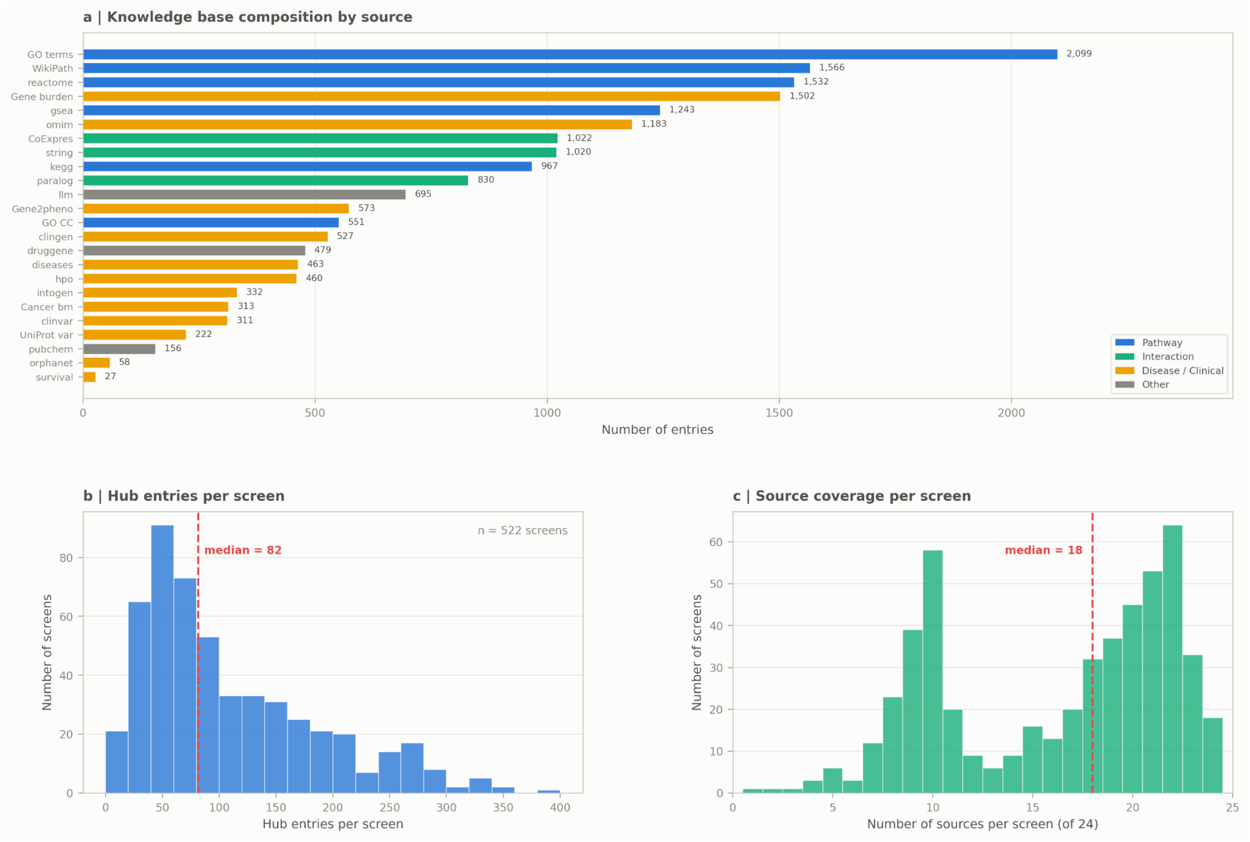


**Supplementary Figure 4 | Resource Hub knowledge-based composition and per-screen evidence coverage. a,** Number of curated entries per knowledge-based source across all hub screens, colored by information type: pathway databases (blue), interaction networks (green), disease and clinical resources (orange) and other (grey). **b,** Distribution of hub entries per screen (n = 522 screens; median = 82, dashed red line). **c,** Number of contributing knowledge-based sources per screen out of 24 total (median = 18).

**Supplementary Table 1 | Gene-level rank changes for autophagy screen 1593**

BioDiscovery and AutoScreen ranks refer to the operational 500-gene output; "—" indicates the gene was not in the output. HIT = YES denotes source-screen validated hits. AutoScreen recovered 26 validated hits in the top 100 versus 20 for BioDiscovery (h@100 +6), and 35 versus 29 at the top 500 (h@500 +6). Sorted by AutoScreen rank.

| Gene | BioDiscovery rank | AutoScreen rank | HIT | Pathway |
|---|---:|---:|:---:|---|
| ATG12 | 57 | 2 | No | LC3 conjugation |
| ATG7 | 56 | 3 | No | LC3 conjugation |
| ATG3 | 50 | 5 | No | LC3 conjugation |
| WIPI2 | 38 | 9 | Yes | Phagophore biogenesis |
| ATG2A | 41 | 10 | Yes | Phagophore biogenesis |
| ATG9A | 43 | 12 | Yes | Phagophore biogenesis |
| RB1CC1 | 28 | 13 | Yes | Autophagy initiation |
| ATG13 | 29 | 14 | Yes | Autophagy initiation |
| ULK1 | 26 | 15 | No | Autophagy initiation |
| BECN1 | 31 | 17 | Yes | PI3KC3 complex |
| PIK3C3 | 32 | 18 | Yes | PI3KC3 complex |

| | | | | |
|---|---|---|---|---|
| ATG101 | 30 | 19 | Yes | Autophagy initiation |
| VMP1 | 185 | 24 | Yes | ER–phagophore contact |
| TMEM41B | 184 | 25 | Yes | ER–phagophore contact |
| NRBF2 | — | 30 | Yes | PI3KC3 complex |
| VPS18 | 128 | 39 | Yes | HOPS complex |
| VPS16 | 127 | 41 | Yes | HOPS complex |
| RAB7A | 112 | 45 | Yes | Vesicle trafficking |
| VPS41 | 131 | 50 | Yes | HOPS complex |
| TRAPPC11 | — | 62 | Yes | TRAPP complex |
| RAB1A | 120 | 67 | Yes | Vesicle trafficking |
| NPC1 | — | 87 | Yes | Cholesterol transport |
| ATP6V1B2 | 89 | 113 | Yes | V-ATPase |
| ATP6V1D | 91 | 267 | Yes | V-ATPase |
| ATP6V1C1 | 90 | 271 | Yes | V-ATPase |
| ATP6V1G1 | 94 | 492 | Yes | V-ATPase |
| MTOR | 1 | — | No | mTORC1 signaling |
| RPTOR | 2 | — | No | mTORC1 signaling |

| | | | | |
|---|---|---|---|---|
| RHEB | 4 | — | No | mTORC1 signaling |
| TSC1 | 14 | — | No | mTORC1 signaling |
| TSC2 | 15 | — | No | mTORC1 signaling |
| AKT1 | 16 | — | No | PI3K–AKT signaling |

**Supplementary Table 2 | Gene-level rank comparison for screen 1161** CTx-DTA is a chimeric toxin combining cholera toxin B subunit (receptor binding via GM1a ganglioside) with diphtheria toxin A (intracellular killing via diphthamide-modified eEF2). BioDiscovery and AutoScreen ranks refer to the operational 500-gene output; "—" indicates the gene was not in the output. HIT = YES denotes source-screen validated hits. AutoScreen recovered 21 validated hits in the top 100 versus 15 for BioDiscovery (h@100 +6), and 35 versus 28 at the top 500 (h@500 +7). DPH3 (rank 8) is not a validated hit (HIT = NO). DPH4 is absent from both ranked outputs. Sorted by AutoScreen rank.

| Gene | BioDiscovery rank | AutoScreen rank | HIT | Pathway |
|---|---|---|---|---|
| UGCG | 1 | 1 | Yes | Ganglioside biosynthesis |
| B3GALT4 | 273 | 2 | Yes | Ganglioside biosynthesis |
| DPH1 | — | 3 | Yes | Diphthamide biosynthesis |

| | | | | |
|---|---|---|---|---|
| ST3GAL5 | 3 | 4 | Yes | Ganglioside biosynthesis |
| B4GALNT1 | 2 | 5 | Yes | Ganglioside biosynthesis |
| DPH2 | — | 6 | Yes | Diphthamide biosynthesis |
| SLC35A2 | 316 | 7 | Yes | Nucleotide-sugar transport |
| DPH3 | — | 8 | No | Diphthamide biosynthesis |
| SPTLC1 | 5 | 9 | Yes | Sphingolipid biosynthesis |
| TM9SF2 | 313 | 10 | Yes | Glycosphingolipid regulation |
| DPH5 | — | 11 | Yes | Diphthamide biosynthesis |
| A4GALT | 4 | 12 | Yes | Glycosphingolipid biosynthesis |
| SPTLC2 | 6 | 13 | Yes | Sphingolipid biosynthesis |

| | | | | |
|---|---|---|---|---|
| CERS2 | 9 | 14 | Yes | Ceramide synthesis |
| DPH6 | — | 15 | Yes | Diphthamide biosynthesis |
| CERS5 | 10 | 17 | Yes | Ceramide synthesis |
| CERS6 | 11 | 18 | Yes | Ceramide synthesis |
| DPH7 | — | 19 | Yes | Diphthamide biosynthesis |
| ATP6V1B2 | 127 | 27 | Yes | V-ATPase |
| ATP6V1E1 | 130 | 28 | Yes | V-ATPase |
| ATP6V1A | 126 | 33 | Yes | V-ATPase |
| ATP6V1C1 | 128 | 46 | Yes | V-ATPase |
| ATP6V1G1 | 132 | 77 | Yes | V-ATPase |
| VPS29 | 75 | 104 | Yes | Retromer |
| VPS53 | 94 | 108 | Yes | GARP complex |
| VPS54 | 95 | 110 | Yes | GARP complex |
| TRAPPC8 | 90 | 174 | Yes | TRAPP complex |
| SEL1L | 33 | 255 | Yes | ERAD |
| SYVN1 | 34 | 256 | Yes | ERAD |
| TRAPPC11 | 91 | 296 | Yes | TRAPP complex |
